# Chemical modification of human decellularized extracellular matrix for incorporation into phototunable hybrid-hydrogel models of tissue fibrosis

**DOI:** 10.1101/2022.10.11.511796

**Authors:** Rukshika S. Hewawasam, Predrag Šerbedžija, Rachel Blomberg, Chelsea M. Magin

**Author notes:** Corresponding Author: Chelsea M. Magin, PhD, 2115 N Scranton Street, Suite 3010, Aurora, CO 80045.

## Abstract

Tissue fibrosis remains a serious health condition with high morbidity and mortality rates. There is a critical need to engineer model systems that better recapitulate the spatial and temporal changes in the fibrotic extracellular microenvironment and enable study of the cellular and molecular alterations that occur during pathogenesis. Here, we present a process for chemically modifying human decellularized extracellular matrix (dECM) and incorporating it into a dynamically tunable hybrid-hydrogel system containing a poly(ethylene glycol)-alpha methacrylate (PEGαMA) backbone. Following modification and characterization, an off-stoichiometry thiol-ene Michael addition reaction resulted in hybrid-hydrogels with mechanical properties that could be tuned to recapitulate many healthy tissue types. Next, photoinitiated, free-radical homopolymerization of excess α-methacrylates increased crosslinking density and hybrid-hydrogel elastic modulus to mimic a fibrotic microenvironment. The incorporation of dECM into the PEGαMAhydrogel decreased the elastic modulus and, relative to fully synthetic hydrogels, increased the swelling ratio, the average molecular weight between crosslinks, and the mesh size of hybrid-hydrogel networks. These changes were proportional to the amount of dECM incorporated into the network. Dynamic stiffening increased the elastic modulus and decreased the swelling ratio, average molecular weight between crosslinks, and the mesh size of hybrid-hydrogels, as expected. Stiffening also activated human fibroblasts, as measured by increases in average cellular aspect ratio (1.59 ± 0.02 to 2.98 ± 0.20) and expression of α-smooth muscle actin (αSMA). Fibroblasts expressing αSMA increased from 24.4% to 51.8% upon dynamic stiffening, demonstrating that hybrid-hydrogels containing human dECM support investigation of dynamic mechanosensing. These results improve our understanding of the biomolecular networks formed within hybrid-hydrogels: this fully human phototunable hybrid-hydrogel system will enable researchers to control and decouple the biochemical changes that occur during fibrotic pathogenesis from the resulting increases in stiffness to study the dynamic cell-matrix interactions that perpetuate fibrotic diseases.

## INTRODUCTION

Progressive tissue fibrosis is a serious health condition that impacts the lungs (idiopathic pulmonary fibrosis, cystic fibrosis), liver (cirrhosis), kidney (renal fibrosis), skin (scleroderma), and other organs.^1^ Fibrosis across all organ types is the result of a failed wound-healing process. Evidence suggests that an insult induces differentiation of fibroblasts into myofibroblasts, which proliferate and deposit excess extracellular matrix (ECM), resulting in fibrotic scarring and tissue stiffening.^1^ This aberrant healing response initiates a positive feedback loop where fibroblasts near the injury activate in response to the stiffened microenvironment, further remodeling the ECM and spreading fibrotic tissue stiffening^2, 3^. Our understanding of the initiation, progression, and potential resolution of fibrosis could be improved by engineering preclinical models that more closely replicate the dynamic nature of this disease.^4^ Here, we present a process for the chemical modification of human decellularized extracellular matrix (dECM) for incorporation into hybrid-hydrogels with a phototunable poly(ethylene glycol)-alpha methacrylate (PEGαMA) backbone to create biomaterial models containing relevant biochemical cues, with dynamic mechanical properties for studying fibroblast-matrix interactions *in vitro*.

Hydrogels, water-swollen polymer networks, are a versatile tool for recapitulating the fibrotic microenvironment and studying specific cell-matrix interactions.^5^ Naturally derived hydrogel biomaterials, including Matrigel,^6–8^ fibrin,^9, 10^ and collagen^11–13^, have been used extensively to study fibrosis in lung, liver, skin, and muscle. Likewise, advancements in tissue-specific decellularization have resulted in dECM hydrogels, which can model disease progression in lung,^14^ liver^15^, and heart.^16^ Although these biomaterials offer environments rich in endogenous biochemical cues, natural polymers that form structures through self-assembly present engineering challenges.^17^ These biomaterials exhibit limited control over mechanical properties and high degradation rates that make it difficult recapitulate the fibrotic microenvironment *in vitro*.^18–21^ One approach to maintain ECM ligands for cellular binding and long-term culture, while enabling mechanical tunability, is chemical modification of natural materials. For example, methacrylation of gelatin and hyaluronic acid resulted in scaffolds that maintained encapsulated valvular interstitial cells (myofibroblast precursors from the heart valve leaflet) for several weeks in culture in a quiescent state.^22, 23^ To decouple the changes in ECM composition from subsequent mechanical changes, Nizamoglu et al. reported a strategy for tailoring the mechanical properties of human lung dECM hydrogels using ruthenium/sodium persulfate-initiated di-tyrosine crosslinking in a way that does not alter the chemical composition.^24^ Similarly, Petrou et al. and Saleh et al. engineered hybrid-hydrogel systems that employed a phototunable PEGαMA backbone as a blank slate for the addition of healthy porcine lung dECM or healthy and fibrotic murine lung dECM, respectively, and showed that dynamic stiffening *in situ* increased fibroblast activation in real-time.^25,26^

Here we build on this work by chemically modifying human dECM for incorporation into a dual-stage polymerization hybrid-hydrogel system and characterizing the resulting hydrogel networks. The dual-stage polymerization technique includes two cytocompatible steps to achieve dynamic stiffening: 1) off stoichiometry thiol-ene Michael addition via step-growth polymerization and 2) homopolymerization of remaining αMA moieties via photopolymerization. Chemical modification of human dECM to convert primary amines to thiols enabled this naturally derived component that enhances cellular adhesion to be incorporated into the dynamic culture system. The incorporation of human dECM increased the molecular weight between crosslinks, swelling, and mesh size of the hybrid-hydrogels compared to synthetic controls, resulting in decreases in bulk elastic modulus measurements directly proportional to the amount of dECM added to the biomaterial system. The dual-stage polymerization provided controlled tunability of biomaterial mechanical properties with average elastic modulus values that matched healthy (4.8 ± 0.19 kPa) and fibrotic (13.89 ± 0.36 kPa) human tissues, pre- and post-stiffening, respectively. This dynamic increase in biomaterial modulus initiated changes in spreading and alpha-smooth muscle actin (αSMA) protein expression consistent with fibroblast activation. These findings enhance our understanding of the biomolecular networks formed within hybrid-hydrogels and how to employ these biomaterial systems to dynamically study the cellular and molecular mechanisms underlying fibrosis.

## EXPERIMENTAL SECTION

### Materials

All chemicals and reagents were purchased from commercial sources and used without further purification unless otherwise stated. Poly-(ethylene glycol)-hydroxyl (PEG-OH; 8-arm, 10 kg mol^−1^) was purchased from JenKem Technology USA Inc. and lyophilized before use. Ethyl 2-(bromomethyl)acrylate (EBrMA) was purchased from Ambeed, Inc. Sodium hydride (NaH), 2-iminothiolane hydrochloride (Traut’s reagent), 3-(trimethoxysilyl)propyl methacrylate, were purchased from Sigma-Aldrich. Human dECM (HumaMatrix) was provide by and purchased from Humabiologics, Inc.

### Methods

#### Poly(ethylene glycol)-alpha methacrylate (PEGαMA) Synthesis

The reaction was carried out under moisture-free conditions. A flame-dried 250 ml Schlenk flask was charged with a stir bar, 10 kg mol^−1^ PEG-OH (5 g, 0.004 mol of -OH), and 80 ml of anhydrous THF (Sigma-Aldrich). Once PEG-OH was completely dissolved, NaH (0.38 g, 0.015 mol, 3.75 molar excess) was added and the reaction mixture was stirred at room temperature for 30 min. EBrMA (3.68 ml, 0.026 mol, 6.5 molar excess) was added dropwise and the Schlenk flask was wrapped in aluminum foil to protect the reaction from light and the mixture was stirred for 48 h at room temperature. The reaction was quenched with 1 N acetic acid and filtered through celite 545. The filtrate was concentrated under reduced pressure and the polymer was precipitated with diethyl ether (Sigma-Aldrich) at 4°C. The polymer was further purified using dialysis (1 kDa MWCO, Repligen) against 3.5 liters of deionized water with a total of four changes over 72 h at room temperature and lyophilized to obtain a pure white solid powder. The functionalization and the purity of the product were verified by ^1^H NMR. Only PEGαMA with functionalization over 95% by comparison of the αMA alkene end group to the PEG backbone was used in subsequent experiments.

#### PEGαMA Characterization

^1^H NMR spectrum was recorded on a Bruker DPX-400 FT NMR spectrometer (300 MHz) and, to ensure proper relaxation of the functional groups and accurate quantification, 284 number of scans and 2.5 s relaxation delay were modified parameters. NMR spectrum was analyzed as follows: chemical shift δ (ppm) (multiplicity, number of protons, and assignment). The chemical shifts for protons (^1^H) were recorded in parts per million (ppm) relative to a residual solvent. ^1^H NMR (400 MHz, CDCl_3_): d (ppm) 1.23 (t, 6H, CH_3_–), 3.62 (s, 114H, PEG backbone), 4.17–4.21 (t, s, 8H, –CH_2_–C(O)–O–O, –O–CH_2_–C(=CH_2_)–), 5.90 (s, 1H, –C = CH_2_), 6.31 (s, 1H, –C=CH_2_). Yield: 74%. End group functionalization of the final PEGαMA polymer was greater than 99% by comparison of the αMA alkene end group to the PEG backbone. (Figure S1)

#### Human dECM Functionalization

Free primary amines on human dECM (HumaMatrix, Humabiologics, Inc) were converted to thiols to create a clickable dECM crosslinker compatible with a thiol-ene Micheal addition reaction. Primary amine concentration was quantified using a standard ninhydrin assay (Sigma-Aldrich) performed according to the manufacturer’s protocol. Traut’s reagent was reacted with the free amines that naturally occur on human dECM proteins in 3 mM ethylenediaminetetraacetic acid (EDTA, Theromfisher) for 1h at room temperature. The molar excess of Traut’s reagent required to minimize amine content following thiolation was identified by measuring amine content after dECM reaction with 5, 10, 25, 50, 75, and 100 molar excess of Traut’s reagent. Thiolated dECM was purified by dialysis (100-500 Da MWCO, Repligen) against deionized water at room temperature and lyophilized to obtain a solid powder. Free thiol content was quantified using Ellman’s reagent (5,5′-dithiobis(2-nitrobenzoic acid), Sigma-Aldrich) according to the manufacturer’s protocol. Absorbance was measured using BioTek Synergy H1: modular multimode microplate reader, with monochromator-based optics and filter-based optics.

#### Human dECM Characterization

Free primary amine content and free thiol content were quantified pre- and post-thiolation treatment. The concentration of protein within each dECM solution pre- and post-thiolation treatment was measured by absorbance (*λ*=260/280) using BioTek Synergy H1: modular multimode microplate reader using a Take3 Micro-Volume Plate. (Figure S2) The protein molecular weight distribution of dECM fragments before and after thiolation was measured by loading equal concentrations of protein based on absorbance measurements of concentration and then separating the proteins using sodium dodecyl sulfate-polyacrylamide gel electrophoresis (SDS-PAGE). Briefly, electrophoresis was performed on 4–20% Mini-PROTEAN TGX Precast Protein Gels for approximately 120 min at 100 V in a tris borate buffer, per the manufacturer’s protocol. A QuickStain Protein Labeling Kit (Amersham) stained bands for fluorescent imaging and qualitative evaluation.

#### Hybrid-Hydrogel Fabrication

Synthesized PEGαMA was reacted with thiolated human dECM and DTT (Sigma-Aldrich) crosslinkers off-stoichiometry (3:8 thiol to αMA) in a base-catalyzed Micheal addition reaction. The weight percent (wt%) of the PEGαMA polymer backbone and the ratio between dECM and DTT crosslinkers were varied to achieve the desired elastic modulus values before and after a secondary crosslinking reaction. TCEP (tris(2-carboxyethyl)phosphine Sigma-Aldrich) was used as a reducing agent to maximize the free thiol content in human dECM. The TCEP molar excess required for maximum free thiol concentration was determined using an Ellman’s assay (Figure S3). Prior to hybrid-hydrogel fabrication, thiolated human dECM (0.012 mg μl^−1^) was dissolved in 250 mM TCEP and incubated for 1 h at a 20-molar excess to the thiol concentration as determined by Ellman’s assay. Stock solutions of PEGαMA (0.4 mg μl^−1^), DTT (Sigma-Aldrich) (250 mM), and CGRGDS (GL Biochem); a peptide sequence that mimics the cell-adhesive ligands on fibronectin (75 mM) were prepared in 0.3 M, pH 8 HEPES ((4-(2-hydroxyethyl)-1-piperazineethanesulfonic acid), Sigma-Aldrich) solution. Both crosslinkers and CGRGDS were combined and added to the PEGaMA to make the precursor solution. Drops (40 μl) of precursor solution were placed between two hydrophobic glass slides covered with parafilm and kept for 30 min at 37°C to polymerize. Polymerized hydrogels were equilibrated in PBS at room temperature with and without 2.2 mM lithium phenyl-2,4,6-trimethyl-benzoylphosphinate photoinitiator (LAP; 0.06 wt%; Sigma-Aldrich) for overnight. Hydrogels in LAP were exposed to UV light (365 nm,10 mW cm^−2^) for 5 min using an OmniCure Series 2000 UV lamp to obtain stiffened hybrid-hydrogels via homopolymerization of free excess αMA moieties.

### Hybrid-Hydrogel Characterization

#### Rheological Measurement of Hydrogel Mechanical Properties

Rheological analysis was performed using a Discovery HR2 rheometer (TA Instruments) with an 8 mm parallel plate geometry and the Peltier plate set at 37°C. Hybrid-hydrogels were analyzed after each polymerization reaction. The geometry was lowered until the instrument read 0.03 N axial force and the gap distance was noted. The gap distance between the plate and the geometry was adjusted until the storage modulus measurement (G’) plateaued and a percent compression of the specific hydrogel was defined and used thereafter. Samples were subjected to frequency oscillatory strain between 0.1 to 100 rad ^s-1^ at 1%. The elastic modulus (E) was calculated using rubber elastic theory and G’ measurements, which were converted assuming a Poisson’s ratio of 0.5 for bulk measurements of elastic hydrogel polymer networks.^27^ To assess whether elastic moduli between soft and stiffened gels were significantly different, a one-way ANOVA with Tukey’s multiple comparison correction was performed.

#### Measurement of Hydrogel Swelling Ratio

Hydrogels were synthesized as described above. Soft and stiffened synthetic and hybrid-hydrogels (N=6) were swollen in deionized water for 48 h to allow hydrogels to swell to equilibrium. Unsubmerged mass (m) and submerged mass (m’) of the just-synthesized hydrogels and swollen hydrogels were measured using the density kit to obtain Buoyancy-based volumetric measurements: V_r_ (relaxed volume) and V_s_ (swollen volume) respectively. Hexane (density (ρ)= 0.659 g ml^−1^) was used as the non-solvent solution for the density kit. V_r_ and V_s_ were calculated using equation (1). ^28^

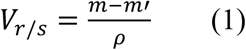

Hydrogels were freeze-dried to remove all solvent and the dried polymer was weighed using the Mettler Toledo MS104TS balance. The dried volume (V_d_) of the polymer was calculated using the PEG polymer density as 1.12 g cm^−3^ and the weight of the dried polymer. The volumetric swelling ratio (Q) was calculated using equation (2). ^28^

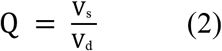

#### Equilibrium Swelling Theory Calculations

Mesh size (ξ) is the average distance between two neighboring network junctions connected by a polymer chain in a hydrogel. In order to estimate mesh size, the average molecular weight between crosslinks in a polymer network (M_c_) was calculated using the Flory-Rehner equation (3), which compares the entropic contribution of mixing polymer and solvent with the elastic energy created as the polymer network swells to incorporate solvent.^29^

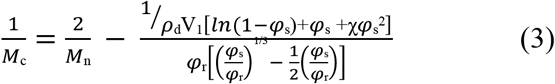

Where M_n_ is the number average molecular weight of an uncrosslinked polymer (10897 g/mol), V_1_ is the molar volume of water (18 mL/mol) and χ is the polymer−solvent interaction parameter (taken to be 0.426 for PEG in water).^30^ φ_s_ is the swollen polymer volume fraction, calculated as V_d_/V_s_ and φ_r_ is the polymer volume fraction in the relaxed state, calculated as V_d_/V_r_. The mesh size was subsequently obtained using the universal version of the Canal-Peppas equation (equation 4).^29^

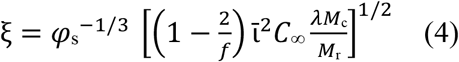

The characteristic ratio (C_∞_) is independent of the length of the chain in a phantom network model and it was taken as 4 for PEG polymers.^28^ The weighted average of polymeric backbone bond length per repeating unit (ῑ) was considered as 0.146 nm.^31^ The molecular weight of the polymer repeating unit (M_r_) was used as 44 g/mol. The number of backbone bonds in the polymer repeating unit (λ) is 3 for PEG polymers. The branching factor of the macromonomers (f) was taken as 8, assuming 100% functionalization with methacrylate groups respectively, and 100% reaction during crosslinking.

#### Estimation of Theoretical Shear Modulus

Shear modulus (G) was calculated according to the Rubberlike elasticity theory using equation (5)^28^. G is the polymer network’s shear modulus, R is the ideal gas constant, and T is the absolute temperature of the system. Frequency of chain-end defects (γ) is considered as 0.5, which would mean that only half of the network chains are connected to two junctions in each condition.

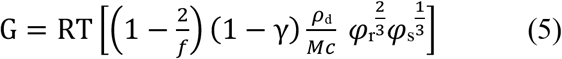

#### Collagen Staining

Hybrid-hydrogels were equilibrated overnight in PBS before being processed for cryosectioning and Picrosirius red staining (Abcam). Hydrogels were submerged in optimal cutting temperature (OCT) compound (Sakura) for 48h at room temperature in a humidified chamber to allow for complete infusion of the compound. Three infused hydrogel samples per condition were stacked in a 15×15×5 mm vinyl cryomold (Sakura) and flash frozen by submersion in liquid nitrogen-cooled 2-methylbutane (Sigma-Aldrich). Frozen samples were removed from the mold and mounted lengthwise onto a Leica CM1850 cryostat such that each cut would take a cross-section of the hydrogel stack. 10-μm sections were acquired from regions throughout the hydrogel stack. Frozen slides were equilibrated to room temperature in deionized water (5-10 min). Then slides were submerged in Picrosiruis red solution for 1 h. Slides were then briefly dipped into two changes of fresh 0.5% acetic acid to clear any unbound dye. Slides were dehydrated by sequential submersion in 100% ethanol (2× 5 min) and SafeClear II (2× 5 min, Fisher Scientific) before being mounted in Cytoseal 60 (ThermoFisher) under a 24 × 50 mm coverslip (Globe Scientific). Imaging was performed on a brightfield microscope (Olympus CKX53 microscope, EP50 camera).

#### Fibroblast Culture

Human pulmonary fibroblasts (ATCC cat. # CCL-151) were passaged at least twice prior to use and incubated in a complete medium consisting of Dulbecco’s Modified Eagle Medium/F-12 (DMEM/F-12) containing 10% fetal bovine serum (FBS, ThermoFisher), 100 units/ml penicillin, 100 units/ml streptomycin, and 0.5 μg/ml amphotericin (Life Sciences) at 37°C and 5% CO_2_. The culture medium was changed every two to three days and the cells were grown to approximately 80% confluency before passaging or seeding onto hybrid-hydrogels.

#### Fibroblast Activation Experiments

Fibroblast activation experiments were conducted under aseptic conditions. Hybrid-hydrogels were fabricated using 15 wt% PEGαMA and a 90:10 DTT:dECM crosslinker ratio as described above. All hydrogel components were dissolved in sterile pH 8 HEPES. Cell culture samples were prepared by silanating 18-mm glass coverslips (Fisher Scientific) with 3-(trimethoxysilyl)propyl methacrylate using a liquid deposition technique.^32^ 90-μl droplets of hybrid-hydrogel precursor solution were placed between each silanted coverslip and a hydrophobic glass slide treated with SigmaCote (Sigma-Aldrich) according to manufacturer instructions. Hybrid-hydrogels were allowed to polymerize for 30 min at 37°C and were equilibrated in sterile PBS for 3h at room temperature. PBS was replaced with complete medium and hydrogels were incubated at 37°C overnight prior to experiments. Fibroblasts were seeded onto hybrid-hydrogels at a density of 20,000 cells/cm^2^ for immunocytochemistry and incubated at 37°C and 5% CO_2_ overnight. The following day (Day 1), the serum content of the culture medium was reduced to 1%. The medium was changed on Day 3 and Day 5. On Day 7, cells on soft hydrogels were processed for immunocytochemistry as described below. To obtain stiffened hybrid-hydrogels, the cell culture medium was supplemented with a 2.2 mM LAP photoinitiator on Day 6 and hydrogels were exposed to UV light (365 nm, 10 mW cm^−2^) for 5 min using an OmniCure Series 2000 UV lamp (Lumen Dynamics). LAP was removed after a 45-min incubation at 37°C and the cell culture medium was replaced to allow for additional incubation at 37 °C and 5% CO_2_ for two more days. On Day 9 cells on stiffened hydrogels were processed for immunocytochemistry.

#### Immunostaining

Unless otherwise specified, all steps were performed at room temperature. Cells were fixed and permeabilized with 0.5 wt% Triton X-100 (0.1%; Fisher BioReagents) in 4% v/v paraformaldehyde in PBS for 10 min. Excess paraformaldehyde was quenched with 100 mM glycine (Sigma-Aldrich) in PBS for 15 min and cells were rinsed with PBS. Non-specific binding sites were blocked with 5% bovine serum albumin (BSA, Sigma-Aldrich) in PBS for 1 h. Mouse anti-human α-smooth muscle actin (αSMA) monoclonal antibody (ThermoFisher Scientific) was diluted 1:250 in 3% BSA/0.1% tween-20 (Fisher Scientific) and incubated for 1 h. Samples were rinsed three times with PBS and incubated in 1:250 goat anti-mouse AlexaFluor555 (ThermoFisher Scientific) secondary antibody in 3% BSA/0.1% tween-20 supplemented with Actin 488 Ready Probes (ThermoFisher Scientific) for 1 h. Nuclei were counter-stained with 0.5 μg/ml DAPI in PBS for 10 min and samples were rinsed three times with deionized water. Cell-seeded hydrogels were mounted onto microscopy slides face up and preserved for imaging with Prolong Gold Antifade (Life Sciences) under a 22-mm coverslip. All microscopy was performed on a BX-63 upright epifluorescent microscope (Olympus). Five fields of view were randomly selected on each hydrogel and imaged with a 10x objective.

#### Image Analysis

Image analysis was performed with Fiji (ImageJ) software (NIH). Cell nuclei were counted on the DAPI channel by establishing a signal threshold, performing a watershed operation, and analyzing particles. Myofibroblasts were counted on the αSMA channel by establishing a signal threshold and analyzing for particles with a circularity less than 0.5, indicating a more spread phenotype as a surrogate for stress fiber formation.

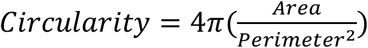

To calculate the percentage of activated cells per sample, the myofibroblast count was divided by the total number of cell nuclei. Cell spreading was quantified on the f-actin channel by establishing a signal threshold and analyzing particles using the fit ellipse function. The aspect ratio was calculated by taking the ratio of the major axis to the minor axis of each ellipse generated by this function. All fibroblast activation experiments are presented as n=6 technical replicates, where each replicate value is the average of five representative fields of view per sample.

## RESULTS AND DISCUSSION

### Chemical Modification of Human Decellularized Extracellular Matrix

During fibrotic progression, the biochemical composition, organization, and biomechanics of the ECM change dramatically^33^. Increased ECM synthesis and remodeling leads to changes in the expression of core matrisome and matrisome-associated proteins and proteoglycans. Excessive accumulation of fibrillar collagen occurs due to an imbalance in collagen synthesis and assembly relative to degradation, which results in tissue stiffening in nearly every major organ.^34^ Healthy liver is typically soft (<6 kPa) with low amounts of fibrillar collagens, such as collagen I.^35^ Collagen mass was shown to increase seven-fold in severely fibrotic livers according to hydroxyproline analysis^36^ and histology.^37^ Subsequently, liver stiffness increased to pathologic levels (>20 kPa); this increase in stiffness was correlated with increased portal pressure, an indicator of cirrhosis.^35^ Analyses of fibrotic human heart tissues revealed increases in collagens I and IV, laminin-***γ***1, fibrillin-1, and periostin. Normal heart tissue has an elastic modulus of 10 to 15 kPa, while fibrotic tissue can be anywhere from two to ten times stiffer.^38^ Excessively stiff heart tissue impacts distensibility, impedes pumping, and contributes to diastolic and systolic dysfunction.^38^ Healthy lung tissue is relatively soft, ranging from 1 to 5 kPa and can stiffen above 10 kPa in pulmonary fibrosis.^39^ These large changes in elastic modulus across various organs are all at least partially attributable to increases in fibrillar collagens, which are also accompanied by higher levels of other ECM proteins, including tenascin-C, fibrillin-1, laminins, and vitronectin.^34^In fact, density measurements of decellularized lung tissues, which contain a high proportion of collagens, revealed that fibrotic ECM was more than twice as dense as ECM from nonfibrotic controls.^40, 41^ Since excessive accumulation of collagen-rich ECM with altered biochemical composition is a hallmark of fibrosis and results in increased tissue stiffness, it is crucial for *in vitro* models of fibrosis to both incorporate and decouple the biochemical and dynamic biophysical characteristics of fibrotic ECM to enable research on each contribution.

Here, we recapitulate healthy and fibrotic extracellular microenvironments by chemical functionalization and incorporation of intact human dECM into dynamically tunable PEG-based hybrid-hydrogels. Naturally occurring free primary amines on the human dECM were thiolated using Traut’s reagent (Figure 1A). The average primary amine concentration of untreated human dECM was 0.41 ± 0.02 μmol mg^−1^ as measured by a ninhydrin assay. Previous studies by our group have measured the primary amine concentration for untreated porcine lung dECM (0.18 ± 0.01 μmol mg^−1^), healthy murine lung dECM (0.40 ± 0.03 μmol mg^−1^), and fibrotic murine lung dECM (0.27 ± 0.01 μmol mg^−1^).^25, 26^ These small differences in primary amine concentration by species and disease state can influence the final functionality of the dECM crosslinker and have been accounted for in subsequent hybrid-hydrogel formulations. To identify the best concentration of Traut’s reagent for conversion of primary amines to thiols, increasing molar ratios of Traut’s reagent were reacted with human dECM and the final primary amine concentration was measured by ninhydrin assay. Primary amine concentration decreased as Traut’s reagent molar excess in the thiolation process increased, plateauing at 75-molar excess (Figure 1B). Using a 75-molar excess of Traut’s reagent, the primary amine concentration of human dECM significantly decreased from 0.41 ± 0.02 μmol mg^−1^ to 0.0063 ± 0.001 μmol mg^−1^, showing that Traut’s reagent reacted with the primary amines to introduce thiol functional groups on intact human dECM. The average free thiol concentration was measured using Ellman’s assay pre-and post-thiolation process. Free thiol concentration was increased significantly from 0.019 ± 0.03 to 0.33 ± 0.03 μmol mg^−1^, achieving 94% conversion of primary amines to free thiols (n = 6, p <0.0001, ANOVA) (Figure 1C). Previous studies that performed similar chemical functionalization of dECM showed that the free thiol concentration in porcine lung dECM was approximately 0.2 μmol mg^−1^ after thiolation and that pepsin-digested healthy murine lung dECM was 0.3 μmol mg^−1^, aligning with the human dECM thiol concentrations achieved through the process in this study.^25,26^

**Figure 1.**
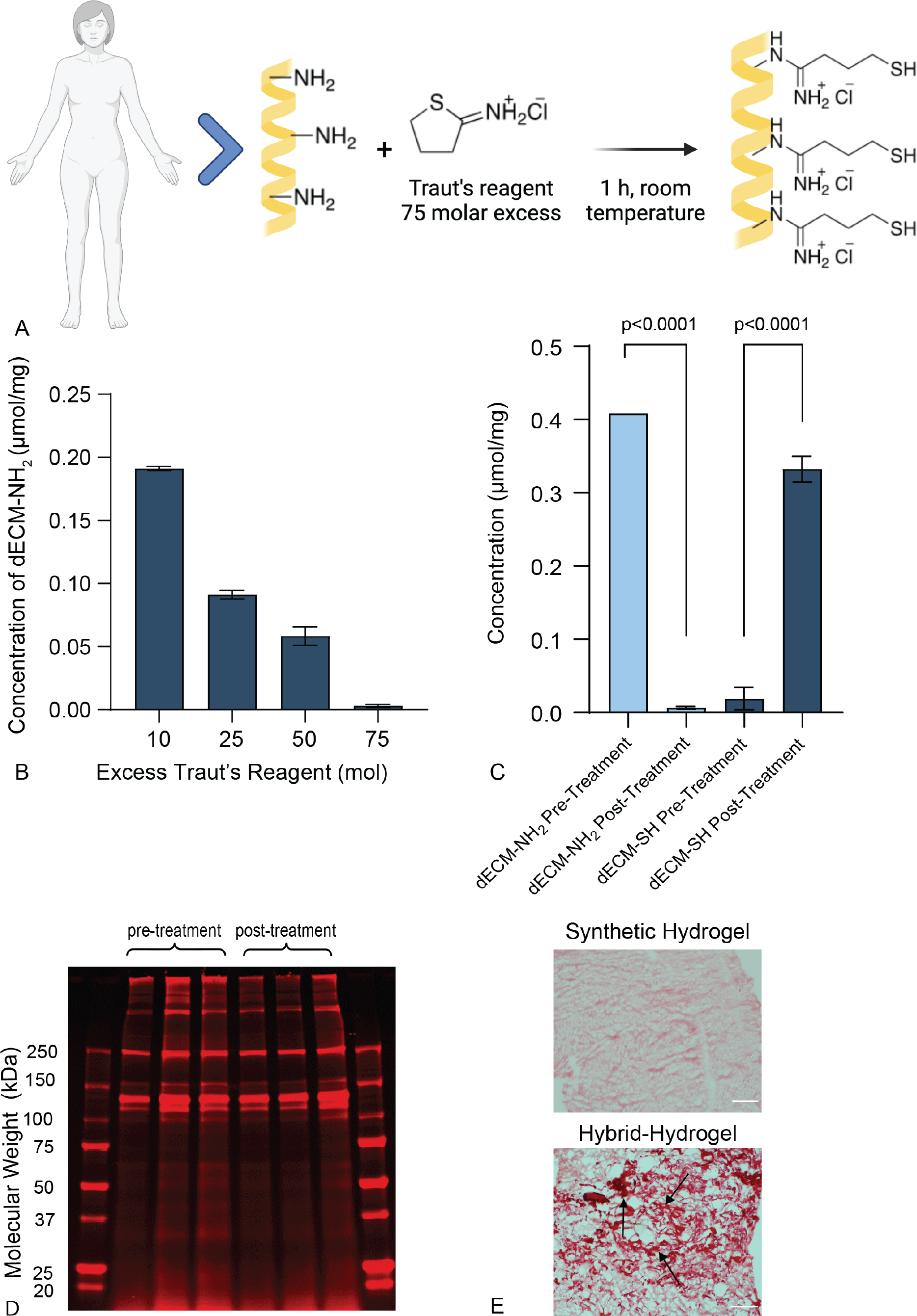
(A) Chemical reaction scheme for human dECM thiolation with Traut’s reagent. (B) Selection of minimum Traut’s reagent molar excess required to convert primary amines to thiols on human dECM. The minimum concentration of primary amine was obtained at 75 molar excess of Traut’s reagent. (N=3, ANOVA) (C) Variation of primary amine and thiol concentration measured in human dECM pre- and post-thiolation process via ninhydrin and Ellman’s assays, respectively. (N=3, Paired t-test). Statistically significant decreases in primary amine concentration and corresponding increases in thiol concentrations demonstrated successful chemical modification of human dECM. (D) SDS-PAGE results verified there was no significant degradation of the protein from the thiolation process. In particular, prominent bands at 250 and 130 kDa showed large protein fragments from human dECM remained intact (n=3). (E) Synthetic and hybrid-hydrogels stained with Picrosirius red to visual dECM distribution. Synthetic hydrogels consisted of 100% DTT crosslinker and hybrid-hydrogels contained crosslinkers that were 10% human dECM and 90% DTT. Black arrows point to collagen strands (dark red) distributed throughout the hybrid-hydrogels.

To verify that the thiolation process did not significantly degrade the proteins that comprise human dECM, protein molecular weight distribution was analyzed pre- and post-thiolation. First, proteins from pre- and post-treatment human dECM were separated by molecular weight using sodium dodecyl sulfate–polyacrylamide gel electrophoresis (SDS-PAGE). Next, total protein was visualized using a Cy5 NHS ester dye. This analysis showed that high molecular weight proteins remained even after the thiolation process, with prominent bands at 250 and 130 kDa, indicating that large protein fragments were not degraded during chemical functionalization and were available for incorporation into hybrid-hydrogels (Figure 1D). Similarly, Burgess et al., showed that protein contents detected by SDS-PAGE in native human lung tissue, decellularized lung, and pepsin-digested lung dECM were consistent, indicating that chemical digestion and mechanical digestion did not degrade the protein content.^42^ This study also assessed dECM using SDS-PAGE and noted prominent bands at 130 kDa and above 250 kDa, in agreement with the results presented here. A Picrosirius red stain was applied to fully synthetic and hybrid-hydrogels to confirm that large collagen fragments were successfully incorporated into these biomaterials^43^. Fully synthetic hydrogels served as negative controls and showed no collagen incorporation by staining light pink with no dark red clusters, while Picrosirius red staining showed homogenously distributed collagen (bright red) within hybrid-hydrogels (Figure 1E).

### Hybrid-Hydrogel Fabrication and Characterization

Following chemical modification, thiolated dECM was incorporated into hybrid-hydrogels featuring a dual-stage polymerization mechanism. Hybrid-hydrogels were comprised of PEGαMA crosslinked with thiolated human dECM and DTT. First, a base-catalyzed thiol-ene Michael addition reaction was carried out off-stoichiometry with a 3:8 ratio between free thiols and α-methacrylate functional groups. Thermodynamically favorable C-S bonds were formed between the α-methacrylate and thiol groups via step-growth polymerization, resulting in soft hybrid-hydrogels (Figure 2A1). Next, excess α-methacrylate groups were subjected to free-radical homopolymerization to achieve dynamic stiffening, increasing the elastic moduli of hybrid-hydrogel samples (Figure 2A2). Both the molar ratio of thiolated human dECM to DTT crosslinkers and the weight percent of PEGαMA were varied to obtain a hydrogel formulation that could exhibit a dynamic stiffening that recapitulates the transition from healthy to fibrotic human tissue.

**Figure 2.**
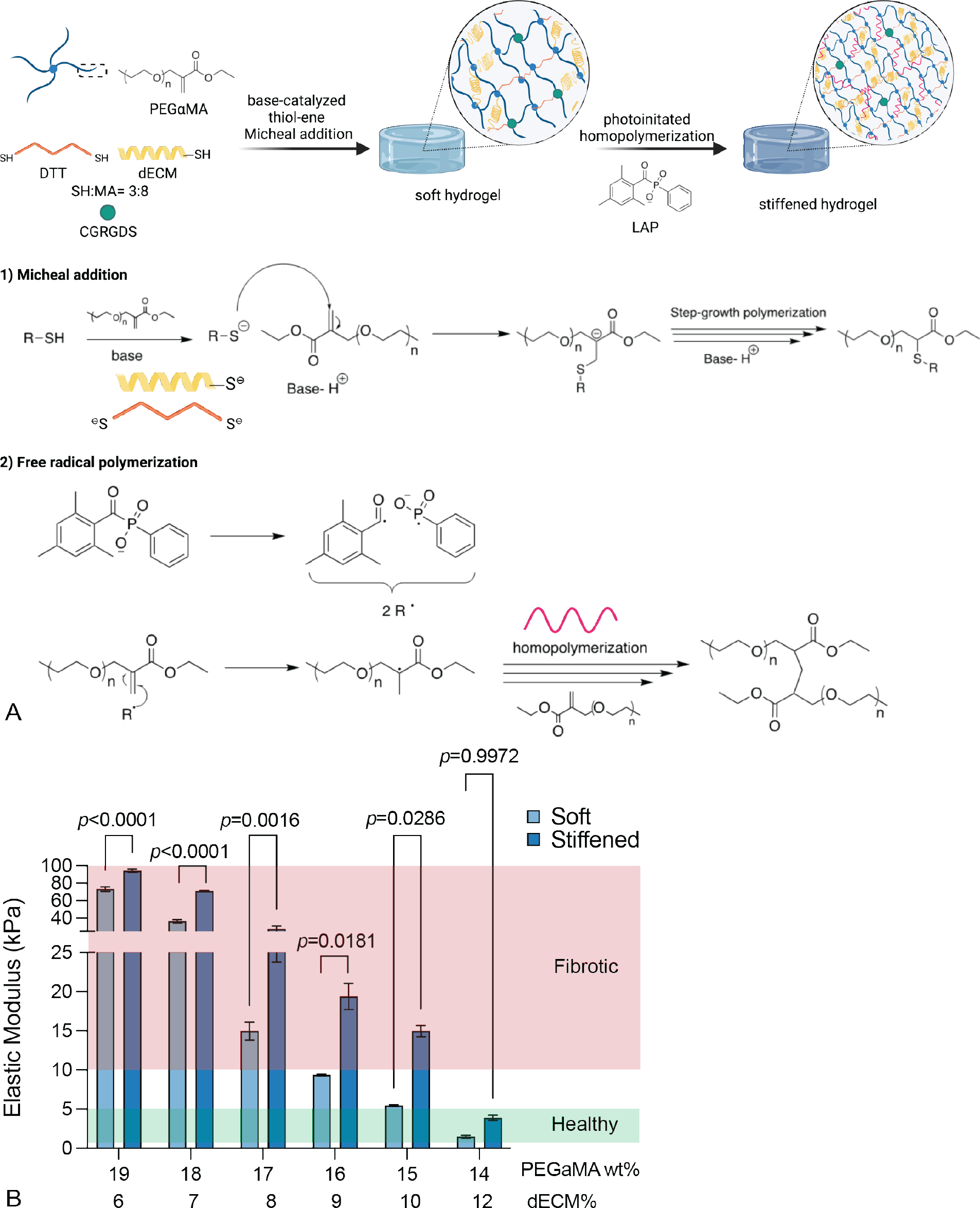
(A) 1. Schematic of the dual-stage polymerization reaction that combined PEGαMA and the chemically modified human dECM crosslinker with DTT and CGRGDS to enable spatiotemporal control over stiffening. Thiol-ene Michael reactions are base-catalyzed reactions where the proton abstraction by the base generates a thiolate anion and a conjugate acid. The potent nucleophile thiolate anion attacks the electrophilic β-carbon of the acrylate, forming the intermediate carbon-centered anion. This anion picks the proton from the conjugate acid and regenerates the base. Subsequently, the anionic propagation undergoes in a stepwise manner. 2. Excess αMA groups in soft hydrogels were subjected to free-radical homopolymerization using LAP as the photoinitiator. UV radiation cleaves LAP homolytically, producing two radicals. Initiator radicals attack the αMA and form a sigma bond between polymer chains increasing the crosslink density and stiffening the hydrogel. (B) The elastic modulus of soft and stiffened hybrid-hydrogels decreased with increasing dECM content ranging from 12 mol% to 6 mol% dECM and with reducing PEGαMA wt% from 19% to 14%. (n = 3, paired t-test)

Rheological studies were carried out to quantify the shear modulus (G) of hybrid-hydrogels containing varying ratios of human dECM:DTT and PEGαMA weight percents. G was converted to elastic modulus E assuming a fully elastic polymer network using a Poisson’s ratio of 0.5 according to the rubber elasticity theory.^44^ For both soft and stiffened hydrogels, elastic moduli decreased with increasing dECM content and decreasing PEGαMA weight percent as expected and as previously demonstrated in other studies.^26^ Soft hybrid-hydrogels ranged from 1.51 ± 0.16 kPa up to 73.41 ± 2.56 kPa and could stiffen from 3.93 ± 0.32 kPa to 94.42 ± 1.98 kPa (Figure 2B). These ranges encompassed the soft and stiffened elastic modulus values of several healthy and fibrotic human tissues. For instance, healthy human liver has a reported elastic modulus of <6 kPa, which can increase to over 20 kPa during fibrotic disease progression.^35^ Healthy human kidney tissues have a healthy modulus of 7.76 ± 5.16 kPa, while fibrotic tissues were measured to be 12.83 ± 9.80 kPa.^45^ Several studies have shown that human healthy lung tissue has an elastic modulus ranging between 1.65 ± 0.2 kPa to 5.6 ± 1.4 kPa, whereas fibrotic human lung tissues have been measured to range from 5.3 ± 1.1 kPa to 16.52 ± 2.25 kPa.^34^ Here, chemically modified human dECM was successfully incorporated into a hybrid-hydrogel system that facilitates spatiotemporal control over precise increases in elastic modulus to recapitulate dynamic increases in human tissue stiffness that occur during fibrotic disease progression.

dECM alone can form hydrogels by self-assembly with concentration-dependent mechanical properties.^46^ Pouliot, et al showed that the shear modulus of human lung dECM hydrogels could range from G=15 to 60 kPa, which was significantly softer than healthy lung tissue.^46^ While naturally derived hydrogels allow for the study of cell-matrix interactions, these systems are not ideal for studying mechanosensing due to the relatively soft, static modulus values achievable with self-assembly. Similarly, using lung dECM, de Hilster et. al. demonstrated that dECM hydrogels formed from fibrotic dECM were stiffer than those formed from healthy dECM; however, these hydrogels were significantly softer than the tissues the dECM was originally derived from, and thus failed to fully recapitulate the physiological microenvironment.^47^ To overcome this limitation, photocrosslinkable methacrylated porcine liver ECM was used to design hydrogels that could achieve stiffnesses as high as 162 kPa, demonstrating the utility of chemically modifying dECM to achieve better control over mechanical properites.^15^

In this report, human dECM was chemically modified for incorporation into hybrid-hydrogels to enable dynamic control over mechanical properties and recapitulate the mechanical properties of healthy and fibrotic human lung tissues. This hybrid-hydrogel system then modeled human pulmonary fibrosis, specifically, the activation of human lung fibroblasts in response to increasing microenvironmental stiffness. Therefore, it is important to note that hybrid-hydrogels formulated with 15 wt% PEGαMA and 10:90 dECM/DTT along with 2 mM CGRGDS exhibited an initial elastic modulus of 4.8 ± 0.19 kPa, recreated the modulus of healthy human lung tissue (1-5 kPa)^48^, and were selected as the starting material for subsequent experiments. The elastic modulus for the stiffened hybrid-hydrogels, obtained after a photoinitiated, free-radical homopolymerization, was 13.89 ± 0.36 kPa, which matched levels of fibrotic lung stiffness (> 10 kPa) (Figure 2B).^34, 48^ This dynamic tunability enabled studies of fibroblast activation in response to increased microenvironmental stiffness within a fully human model of pulmonary fibrosis.

The influence of dECM content on hybrid-hydrogel network formation and subsequent mechanical properties was analyzed and compared to fully synthetic PEGαMA controls. The elastic moduli and swelling ratios for both synthetic and hybrid-hydrogels were measured and used to calculate the average molecular weight between crosslinks, mesh size, and theoretical storage modulus (G’) based on equilibrium swelling theory. The synthetic hydrogel was synthesized using 15 wt% PEGαMA along with 100% DTT. Hybrid-hydrogels were synthesized using 15 wt% PEGαMA with varying ratios of dECM:DTT (2.5%, 5%, 7.5%, and 10%), i.e., increasing amounts of dECM crosslinker. The elastic modulus was directly related to the amount of human dECM present in hybrid-hydrogel. As dECM content increased, the elastic modulus of hybrid-hydrogel samples decreased (Figure 3A). The elastic modulus of soft fully synthetic hydrogels was 23.22 ± 0.25 kPa, while the elastic modulus of soft hybrid-hydrogels containing 10% human dECM was 4.62 ± 0.41 kPa, indicating that the introduction of human dECM significantly lowered the elastic modulus. A similar trend was measured for both soft and stiffened conditions. Conversely, the equilibrium swelling ratio of hybrid-hydrogels was significantly higher than that of synthetic hydrogels, indicating that incorporation of human dECM led to higher water absorption in the hydrogel (Figure 3B). The volumetric swelling ratio increased from 2.07 ± 0.15 for fully synthetic hydrogels to 18.81 ± 0.83 for hybrid hydrogels containing 10% dECM. This phenomenon could be due to the strong hydrogen bonding between protein fragments in dECM and water molecules, which led to higher water molecule retention in the hydrogel network. Moreover, the swelling capacity of soft hybrid-hydrogels was twice that of the stiffened, indicating that free-radical homopolymerization did increase the crosslinking density of the network. A similar pattern of increased swelling and decreased modulus was observed by Yang et al. when increasing the azide-functionalized RGD peptide concentration in PEG-peptide hybrid-hydrogels.^49^ Rizzi, et al. have shown that the elastic moduli and swelling ratios of the protein-*co*-PEG hydrogels could be varied as a function of the stoichiometry of the reactive groups and precursor concentration.^50^ These hybrid-hydrogel networks exhibited higher values for elastic modulus and lower values for swelling ratio when the thiol content on precursor proteins increased. In this study, the same thiol concentration was maintained for each concentration of human dECM. Hence, hybrid-hydrogels with high dECM content showed lower elastic modulus and higher swelling capacity. The same trend of swelling degree and crosslinking density was observed in photoinitiated methacrylated-hyaluronic acid hydrogel networks where high macromer concentration in networks exhibited lower swelling ratios and higher crosslinking densities.^51^ The changes in the swelling ratio were characterized as a function of degradation time in degradable synthetic hydrogel networks, furthermore, and swelling ratios and degradation rates were compared to changing chemistries between copolymerizing macromers.^52^ These studies reinforced that the chemistry and functionality of hydrogel networks directly influence hydrogel mechanical properties.

**Figure 3.**
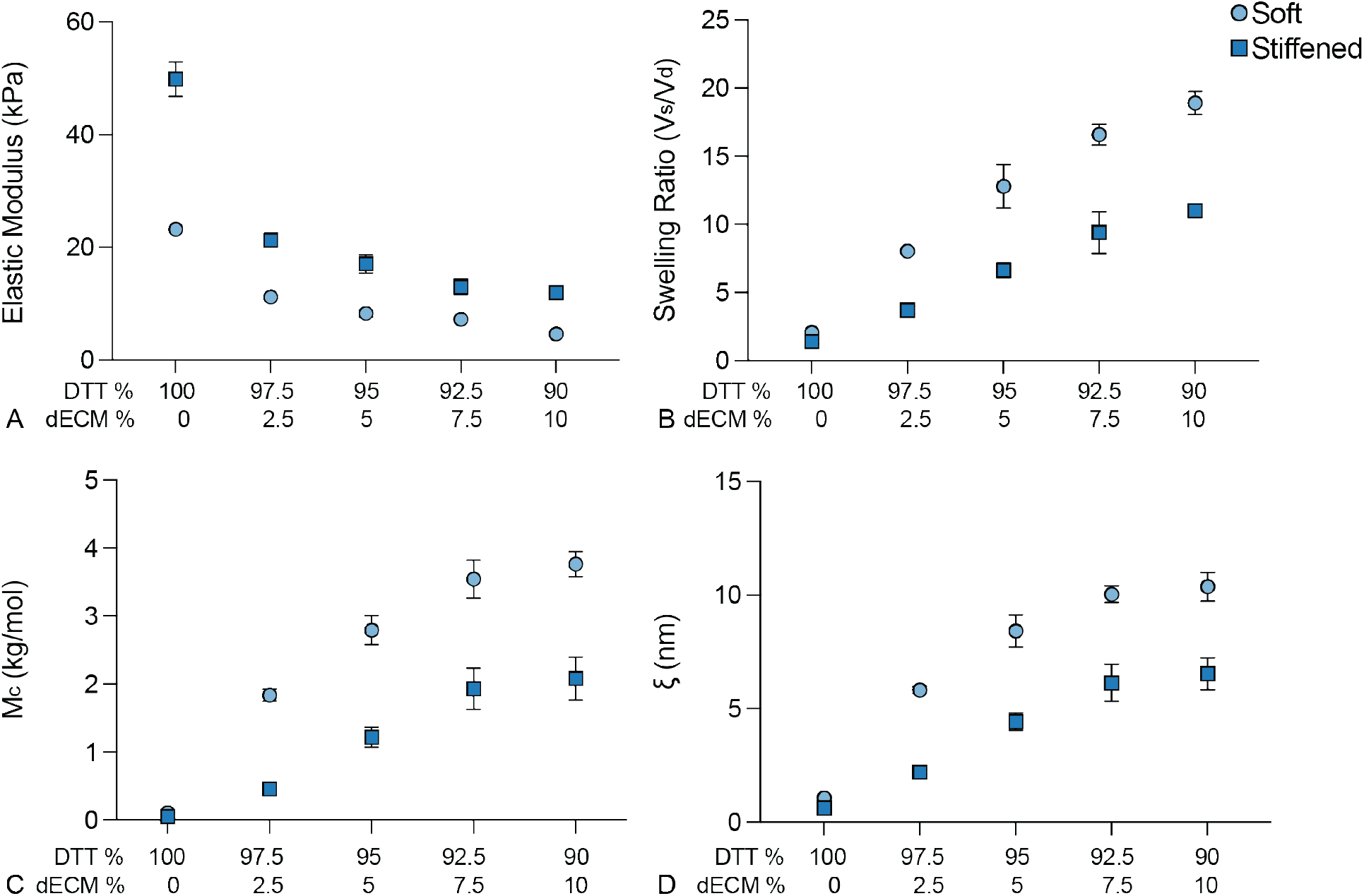
Mechanical properties of hybrid-hydrogels at different human dECM mol% (A) The elastic moduli of soft and stiffened hybrid-hydrogels decreased with increasing dECM content ranging from 0 mol% to 10 mol% dECM (n = 6, error bards, SEM). (B) Volumetric swelling ratio (C) number average molecular weight between crosslinks in a polymer network (M_c_), and (D), mesh size of the network respectively increased with increasing dECM content ranging from 0 mol% to 10 mol% dECM (n = 6, SEM). 100% DTT with 0% dECM; synthetic hydrogel has lower swelling, M_c_ and mesh size compared to the hybrid-hydrogels.

The number average molecular weight between crosslinks (M_c_) was calculated to confirm this observation. M_c_ and mesh size followed the same trend as the swelling ratio, indicating that the incorporation of human dECM increased the distance between the network junctions. The M_c_ was calculated to be 0.11 ± 0.01 kg mol^−1^ for the soft synthetic hydrogel and 3.77 ± 0.18 kg mol^−1^ for the soft hybrid-hydrogel with 10% human dECM. This significant increase in M_c_ was due to both the higher molecular weight of the dECM crosslinker, which exhibited protein fragments up to 250 kDa compared to DTT (154.23 g mol^−1^), and may also have been influenced by the unfolding of the coiled structure of human dECM. Previous studies of PEG-peptide hydrogels used circular dichroism spectrometry to demonstrate that coiled peptide structures influenced entanglements and resulted in a higher degree of swelling.^49^ In natural hydrogels, collagen chains were observed to form new hydrogen bonds with water as well as within the collagen chain itself, thereby increasing the swelling ratio of collagen-based hydrogels.^53^ Similarly, the coiled structure of human dECM used in this hybrid-hydrogel could lead to entanglements and increase hydrogen bonding between water molecules and hydrogel network. This reasoning would predict that, compared to synthetic hydrogels, hybrid-hydrogels should display a higher level of swelling, higher M_c_ and higher mesh size, as measured here. These results were also in agreement with a study that observed increased swelling capacity of poly(acrylic acid) hydrogels with the incorporation of up to 6.7 wt% pectin. Interestingly, increasing the amount of pectin beyond 6.7 wt% led to a decrease in swelling, illustrating that beyond a point, higher crosslinking density will inevitably lead to lower mesh size in the network, reducing the swelling capacity.^54^ These results have implications for future hybrid-hydrogel designs using dECM. In this study, incorporation of up to 10% dECM resulted in increased swelling ratios that appeared to be reaching a plateau (Figure 3B). Increasing the amount of dECM beyond 10% in hybrid-hydrogel formulations may lead to decreases in the swelling ratio. At the ratios studied here, the incorporation of dECM into hybrid-hydrogels appeared to increase hydrogen bond formation within the peptide fragments, contributing to increased swelling and higher mesh size in the network.

The estimated mesh size of soft synthetic hydrogel was calculated to be 10.7 ± 0.20 nm compared to 10.37 ± 0.63 nm for the soft hybrid-hydrogel consisting of 10% human dECM, indicating that the mesh size of the network was significantly increased with the incorporation of human dECM. Since the overall mesh size was higher in hybrid-hydrogels than in fully synthetic hydrogels, these biomaterials could be more permissive to the diffusion of bioactive molecules and migration of cells through the complex network structure, making the use of these hybrid-hydrogels within three-dimensional (3D) models even more physiologically relevant. Also, lower estimated mesh size and M_c_ for the stiffened conditions compared to soft hydrogels were observed again showing crosslink density increased after the photo-initiated homopolymerization of excess αMA moieties.

The theoretical and measured shear moduli of synthetic hydrogels and hybrid-hydrogels containing 10% dECM and synthetic hydrogel containing 100% DTT as the crosslinker were compared to better understand changes in network structure and mechanical properties that occurred when adding dECM into the PEG-based system (Table 1). Synthetic hydrogel theoretical shear moduli were measured to be 4.7 ± 0.2 kPa and 5.98 ± 0.5 kPa for soft and stiffened conditions respectively, whereas measured shear moduli for synthetic hydrogels were 1.31± 0.01 kPa and 4.17 ± 0.1 kPa, for soft and stiffened conditions respectively, giving a greater difference between theoretical and measured shear moduli for synthetic hydrogels than for hybrid-hydrogels in both soft and stiffened conditions. These differences between theoretical and measured shear modulus may be attributed to non-ideal crosslinking behavior.^55, 56^ Poor accessibility of crosslinking sites during gelation, due to steric hinderance or crosslinker molecular weight, could have led to the presence of dangling chains within the polymer network where one end is attached at a crosslinking point and the other end is not attached to the network and thus elastically inactive.^50^ Synthetic hydrogels contain only DTT, a short crosslinker (154 g mol^−1^), which could have led to dangling chains because it was not long enough to react with two other molecules. An increased number of dangling chains lowers the overall crosslinking density, thereby reducing elastic recovery and increasing swelling ratio. For example, in 4 arm star PEG (25 kDa) hydrogels, when the concentration was too low or too high, a short crosslinker could not link the star polymer efficiently. The resultant hydrogel network contained defects such as dangling chains or loops and showed heterogeneity in the viscoelastic measurements.^57^

**Table 1.** Comparison of Theoretical and Measured Shear Moduli (G) of Synthetic and Hybrid-Hydrogels.

|  | Theoretical G<sup>a</sup><br>(kPa) | Measured G<sup>b</sup><br>(kPa) |
| --- | --- | --- |
| Synthetic (soft) | 4.70 ± 0.2 | 1.31 ± 0.01 |
| Hybrid (soft) | 3.26 ± 0.1 | 1.6 ± 0.09 |
| Synthetic (stiffened) | 5.98 ± 0.5 | 4.17 ± 0.10 |
| Hybrid (stiffened) | 5.02 ± 0.6 | 4.64 ± 0.14 |

<sup>a</sup>Theoretical G was calculated by rubberlike elasticity theory using equation (5).

Incorporating human dECM into the PEG hydrogels potentially decreased the number of dangling chains because human dECM was significantly higher in molecular weight (~ 130-250 kDa). Even though a greater extent of dangling chains has been observed to result in a higher swelling ratio^58^, synthetic hydrogels displayed reduced swelling compared to hybrid-hydrogels. This result may be because dECM can have H-bond interactions with water molecules at equilibrium, which creates a stronger contribution to swelling than the dangling chains in the synthetic hydrogel. Kyburz and Anseth have illustrated that a network with a high degree of physical entanglements can display a greater measured modulus than the theoretical modulus.^59^ A reduced difference between theoretical and measured shear moduli for hybrid-hydrogels may be attributed to the physical entanglements of the human dECM in the hydrogel.

Many strategies have emerged recently to functionalize biomacromolecules for incorporation into biomaterials that present tissue-specific cues.^60^ Here, chemically functionalized human dECM was successfully incorporated into hybrid-hydrogels to recapitulate human healthy and fibrotic tissues. The incorporation of human dECM into hybrid-hydrogels not only provided a physiological biochemical component to the biomaterials, but also significantly influenced network properties such as swelling, average molecular weight between crosslinks, and mesh size.

### Cell Activation

Myofibroblasts are critical mediators of fibrotic progression. Previous studies have shown that composition and increases in microenvironmental stiffness can promote the activation of fibroblasts to myofibroblasts, as demonstrated by both the expression of αSMA stress fibers and enhanced cell spreading.^61, 62^ Booth et al. observed a significant increase in αSMA expression on acellular fibrotic human lung tissue compared to normal human lung tissue and demonstrated that fibrotic dECM was sufficient to induce fibroblast activation.^63^ Due to the simultaneous biochemical and mechanical changes that occur during fibrosis, it was not possible in that study to determine which factor was the more potent driver of fibroblast activation. Studies of cellular responses to increased microenvironmental modulus have therefore relied on hydrogels with tunable mechanical properties. For example, a pronounced increase in αSMA expression in cardiac fibroblasts was observed on stiffer hyaluronic acid hydrogels (50 kPa) compared to cells cultured on 8 kPa hyaluronic acid hydrogels that recapitulate healthy myocardium.^64^ However, recent studies using models designed to decouple fibrotic composition from mechanical stress showed that material stiffness is the dominant factor that affects the activation of fibroblasts.^65–67,26^ In particular, Nizamoglu et al. created hydrogels from porcine dECM that could be stiffened by using a ruthenium and sodium persulfate-initiated reaction to crosslink tyrosine moieties on the dECM. Results of fibroblast activation studies using this model showed increased activation induced by stiffening, even when the dECM composition of the hydrogel remained the same^66^. Similarly, murine fibroblasts cultured on PEG-based hybrid-hydrogels consisting of porcine dECM^25^ and murine dECM^26^ showed prominent Col1a1 and αSMA expression on stiffened hydrogels. When dECM derived from fibrotic murine lung was incorporated into hybrid-hydrogels, it only induced modest changes to fibroblast activation relative to the robust changes induced by stiffness.^26^

In this study, human fibroblasts were seeded on 2D soft hybrid-hydrogels and then exposed to dynamic stiffening via dual-stage polymerization. Fibroblast activation was measured by assessing cell morphology based on f-actin expression and myofibroblast phenotype by αSMA expression, the combination of which correlates to high expression of a fibroblast activation genetic program^68^. Immunofluorescence staining for f-actin revealed a significant increase in cellular aspect ratio, the ratio between the major and minor axis of an ellipse fit to the cell area. Increasing aspect ratio indicates increased cellular elongation. On soft hybrid-hydrogels, fibroblasts remained rounded, while fibroblasts on stiffened hybrid-hydrogels began to stretch out protrusions that increased this aspect ratio from 1.59 ± 0.02 to 2.98 ± 0.20 (Figure 4C). Similarly, stiffening resulted in a greater proportion of non-circular, αSMA+ myofibroblasts^68^ from 25.4% on soft hybrid-hydrogels to 51.8% on stiffened samples (Figure 4D). These results demonstrate the ability of this fully human hybrid-hydrogel culture model to facilitate investigation of dynamic fibroblast mechanosensing, a critical component of fibrotic progression.

**Figure 4.**
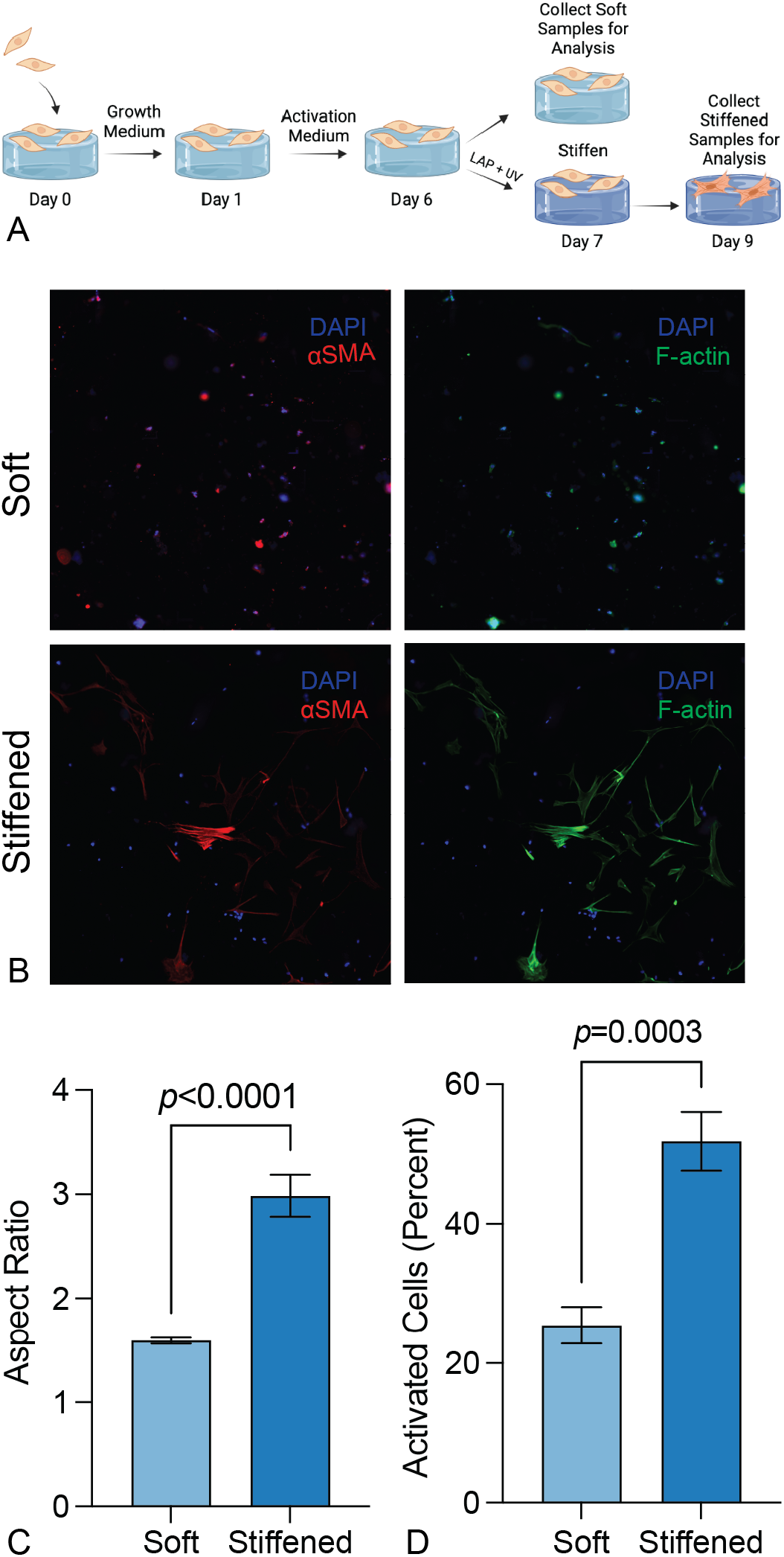
(A) Schematic diagram of the timeline for cell activation experiments. HPFs were cultured on soft hydrogels on day 0. Cells were cultured in 1% FBS media for all conditions. The photoinitiator (LAP) was added to culture media on day 6 for hydrogels to be stiffened and 365 nm UV light at mW cm^−2^ was applied for 5 min at day 7. Samples were collected and analyzed on day 7 and day 9 for soft and stiffened, respectively. (B) Representative images of human pulmonary fibroblasts on soft and stiffened hybrid-hydrogels showed expression of f-actin (green) and αSMA (red) with the DAPI staining (blue). On stiffened conditions, fibroblasts demonstrated increased spreading and more stress fibers morphology compared to fibroblasts cultured on soft hybrid-hydrogels. (C) Quantification of average cellular aspect ratio expressing higher cell spreading on stiffened hybrid-hydrogel. (p< 0.0001, N = 6) (D) Quantification of the percentage of fibroblasts displaying higher αSMA expression on stiffened hybrid-hydrogel, which indicates the fibroblast activation upon stiffening (p=0.0003, N = 6).

## CONCLUSION

A dynamically tunable hybrid-hydrogel system containing chemically modified human dECM was engineered to study fibroblast-matrix interactions *in vitro*. By incorporating intact, tissue-derived dECM, key biochemical properties of human tissue were recapitulated while taking advantage of dual-stage photopolymerization techniques to mimic the dynamic increase in microenvironmental stiffness observed during fibrotic progression in human organs. The incorporation of dECM altered key network properties of the hydrogels, including increasing the swelling ratio, average molecular weight between crosslinks, and mesh size, while decreasing the average shear modulus of the resulting hybrid-hydrogels. Focusing on the lung as a model system, fibroblast activation was measured in response to stiffening on human hybrid-hydrogels and was shown to increase with increasing elastic modulus. This hybrid-hydrogel system could support encapsulation of cells in 3D to investigate cellular responses to dynamic stiffening in an environment that is both biochemically relevant and mechanical tunable. Hybrid-hydrogels could also be generated with dECM sourced from specific tissues to provide the most physiologically relevant environment for a variety of cell types, improving the study of cellular and molecular mechanisms underlying fibrotic disease initiation and progression.

## AUTHOR CONTRIBUTIONS

RH, PS, RB, and CMM conceived the research plan. RH, PS, and RB carried out the experiments. All authors wrote, reviewed, and edited the manuscript.

## ACKNOWLEDGEMENTS

This work was supported by funding from the National Heart, Lung, and Blood Institute of the National Institutes of Health (NIH) under awards R01 HL153096 (CMM, PS, and RB), and T32 HL 07085 (RB); the National Cancer Institute of the NIH under award R21 CA252172 (CMM and RB); the National Science Foundation under award 1941401 (CMM and RH); the Department of the Army under award W81XWH-20-1-0037 (CMM).

