## Supplementary Materials for "Chemical modification of human decellularized extracellular matrix for incorporation into phototunable hybrid-hydrogel models of tissue fibrosis"

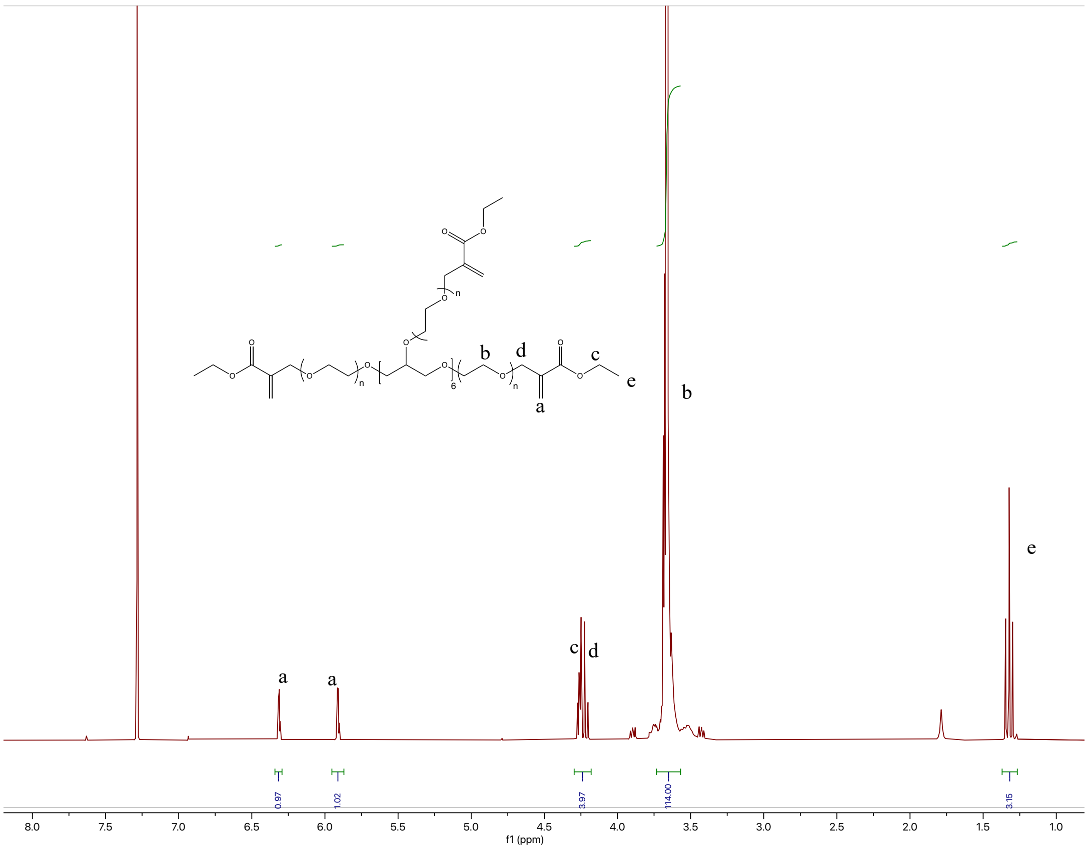


Figure S1. PEGαMA [1]H NMR (300 MHz, CDCl3): δ (ppm) 1.36 (t, 3H, $CH_3$–), 3.71 (s, 114H, PEG $CH_2$-$CH_2$), 4.29 (t, s, 4H, –$CH_2$–C(O)–O–O, –O–$CH_2$–C(=$CH_2$)–), 5.93 (q, 1H, –C=$CH_2$), 6.34 (q, 1H, –C=$CH_2$). End group functionalization of the final PEGαMA polymer was greater than 99% by comparison of the αMA alkene end group to the PEG backbone

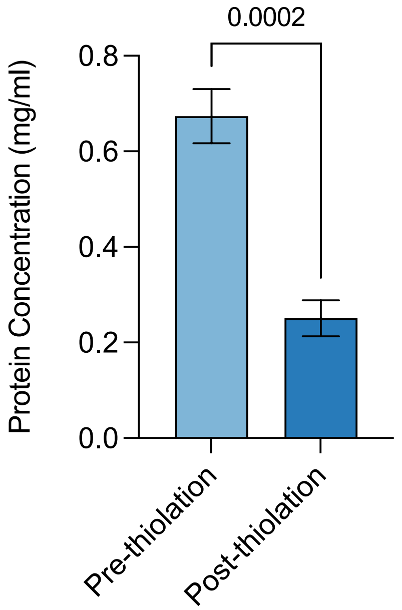


Figure S2. Total protein concentration measured by absorbance ($\lambda$=260/280). (N=5, Paired t-test). Protein concentration as measured by absorbance significantly decreased after thiolation.

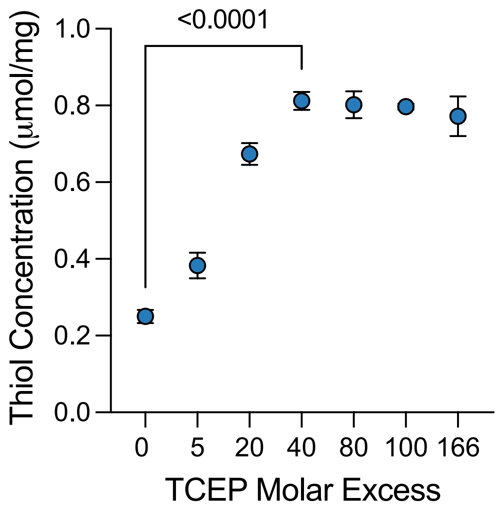


Figure S3. Human dECM was treated with excess TCEP ranging from 0 to 166 molar excess to identify the quantity required to obtained maximum thiol concentration via Ellman's assay. (N=3, ANOVA).
